# Response to “No evidence that transmissible cancer has shifted from emergence to endemism in Tasmanian devils”

**DOI:** 10.1101/2025.07.14.664746

**Authors:** Andrew Storfer, Mark J. Margres, Christopher P. Kozakiewicz, Rodrigo Hamede, Manuel Ruiz-Aravena, David G. Hamilton, Sebastian Comte, Tanja Stadler, Adam Leaché, Hamish McCallum, Menna Jones, Paul A. Hohenlohe

## Abstract

Herein, we rebut the critique of Patton et al. (2020), entitled, “No evidence that a transmissible cancer has shifted from emergence to endemism”, by Stammnitz et al. (2024). First and foremost, the authors do not conduct any phylogenetic or epidemiological analyses to rebut the inferences from the main results of the Patton et al. (2020) article, rendering the title of their rebuttal without evidence or merit. Additionally, Stammnitz et al. (2024) present a phylogenetic tree based on only 32 copy number variants (not typically used in phylogenetic analyses and evolve in a completely different way than DNA sequences) to “rebut” our tree that was inferred from 436.1 kb of sequence data and nearly two orders of magnitude more parsimony-informative sites (2520 SNPs). As such it is not surprising that their phylogeny did not have a similar branching pattern to ours, given that support for each branch of their tree was weak and the essentially formed a polytomy. That is, one could rotate their resulting tree in any direction and by nature, it would not match ours. While the authors are correct that we used suboptimal filtering of our raw whole genome sequencing data, re-analyses of the data with 30X coverage, as suggested, resulted in a mutation rate similar to that reported in Stammnitz et al. (2024). Most importantly, when we re-analyzed our data, as well as Stammnitz et al.’s own data, the results of the Patton et al. (2020) article are supported with both datasets. That is, the effective transmission rate of DFTD has transitioned over time to approach one, suggesting endemism; and, the spread of DFTD is rapid and omnidirectional despite the observed east-to-west wave of spread. Overall, Stammnitz et al. (2024) not only fail to provide evidence to contradict the findings of Patton et al. (2020), but rather help support the results with their own data.

## 1. Introduction

Stammnitz et al. [1] claim that there is “No evidence that transmissible cancer has shifted from emergence to endemism in Tasmanian devils,” which is presented as a contradictory argument to Patton et al. [2]. Much of their criticism addresses the limitations of our dataset for genome-wide mutation detection in samples of devil facial tumor disease (DFTD), a fatal, transmissible cancer. However, this was not the goal of our phylodynamic analyses [in 2], which were to estimate epidemiological parameters of DFTD, including the directionality of spread and temporal variation in transmission rates. Notably, the authors do not present epidemiological evidence contrary to that presented in Patton et al. [2], which shows a gradual decline in the effective reproduction number (*R*_*E*_, used as a proxy for transmission rate) of DFTD from ∼3.5 early in the epizootic to 1 at present, suggesting a shift from emergence to endemism (supplemental data located: science.sciencemag.org/content/370/6522/eabb9772/suppl/DC1; scripts for data analyses can be found at: https://zenodo.org/records/14207466). Further, Stammnitz et al. [1] do not acknowledge an increasing and independent body of published work supporting these results. That is, despite the ubiquity of DFTD across the Tasmanian devil’s geographic range and a near 100% case fatality rate: 1) no population has become extirpated [3]; 2) some populations may be recovering [4]; 3) there are an increasing number of spontaneous tumor regressions in the wild [5]; 4) there is ample evidence for genomic responses to selection from DFTD across the devil genome [6–8]; 5) increasing evidence for sufficient genomic variation to respond evolutionarily to the strong selection imposed by DFTD [9]; 6) models parameterized using more than 25 years of field mark-recapture data show DFTD extinction or devil-DFTD coexistence in the majority of scenarios [10]. Moreover, a recently published genome-wide association study of the devil-DFTD genomic interaction indicates strong evidence for coevolution at the genomic level [11]. In summary, multiple independent lines of evidence using models, field studies, genomic data and (co)evolutionary analyses utilizing data generated from decades-long studies of wild Tasmanian devil populations, taken together, also support a declining threat of DFTD to devil persistence.

Stammnitz et al. [1] correctly point out that we could have provided more detail regarding our filtering of genomic data used to call variants necessary for the original phylodynamic analyses presented in Patton et al. [2]. We also agree with their argument that our original data filtering procedure was not optimal, as it resulted in a surplus of singletons (i.e., mutations found in single tumor isolates) that inflated our initial estimates of mutation rates. Consequently, to explore the robustness of our original findings, we employed stringent filtering to re-analyze our data to identify higher-confidence single nucleotide polymorphisms (SNPs) in tumor genomes. Details of the filtering procedure are described below in the Materials and Methods, but we notably include a minimum mean sequencing depth coverage of 30x for calling variants (as recommended in Stammnitz et al. [1]). As in Patton et al. [2], we screened the resulting conservative set of somatic mutations for a strong, positive clock-like evolutionary signal (as recommended in [12]). Re-filtering ultimately led to the retention of 3,591 SNPs (mean sequencing depth ± 1SD = 50.75 ± 18.42; Table S1) among the original 54 tumor samples (Table S2) that were subsequently re-analyzed using the Birth-death skyline model [13] routinely used for phylodynamic analyses. An additional benefit of filtering sequencing data for a clock-like evolutionary signal is the removal of loci with temporal shifts in mutation rates that can lead to inaccurate phylogenetic reconstruction (note all raw data and scripts for reanalysis of our data and Stammnitz et al’s [1,14] data have been uploaded to DOI: 10.5281/zenodo.14207593).

Here, we show that the results reported in Patton et al. [2] are robust, with an *R*_*E*_ [12,13] of DFTD early in the epizootic estimated to be ∼3.5, followed by a gradual decline to just below 1 (replacement) among extant tumor lineages (Fig. 1a). These results support our initial conclusions that DFTD transmission has shifted from emergence (and an initial exponential increase in disease prevalence) to endemism at present (Fig 1a). We emphasize here that we *also* analyzed the supporting data provided in Stammnitz et al. [1,14] (see also Materials and Methods; Table S3, S4) and find the same trend, irrespective of whether tetraploid samples are included (Fig. 1b) or excluded (Fig. 1c). Although the timing of peak transmission and the subsequent decline in *R*_*E*_ differ slightly across datasets from [1,2,14], the overall conclusion from these analyses remains the same: DFTD appears to be transitioning towards endemism.

**Figure 1.**
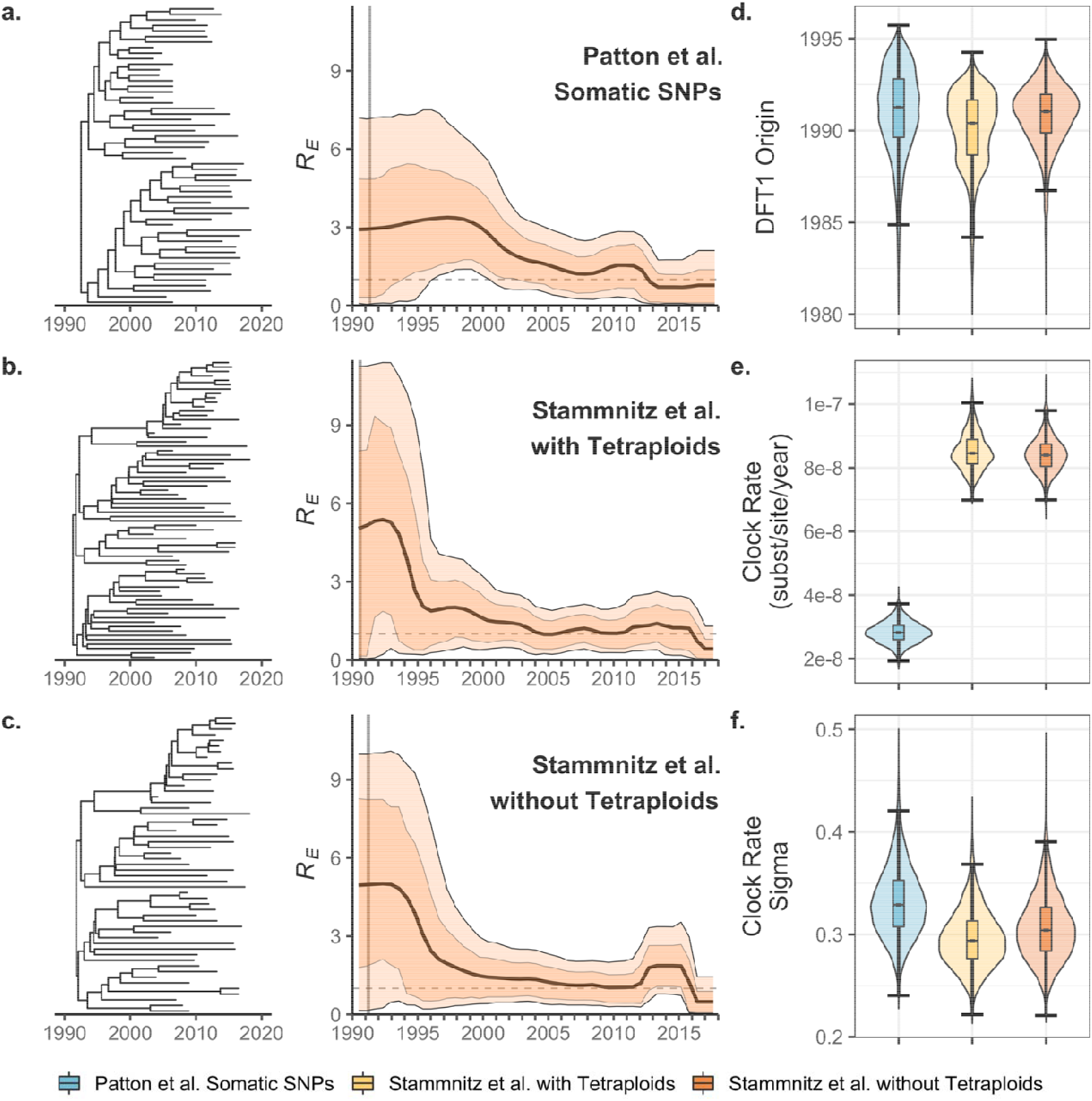
Independent datasets suggest DFTD/DFT1 is transitioning from emergence to endemism. Maximum clade consensus trees (left) and Birth-death skylines of the effective reproduction number (R_E_) through time as estimated using the re-filtered data of Patton et al. [2], Stammnitz et al. [14], either including (**b**.) or excluding (**c**.) tetraploid samples. Median estimates of *R*_*E*_ are plotted as bold lines, and the 80% and 95% highest posterior density of estimates are shown as the inner (darker) and outer (lighter) bands respectively. Vertical grey line indicates the median estimate of the origin of DFTD. Posterior distributions of the: (**d**.) origin of DFTD, (**e**.) clock rate (substitutions/site/year), (**f**.) clock rate variation (i.e., sigma), the variance of the lognormal distribution from which rates are drawn.

Our re-analysis infers the date of origin of DFTD to be 1991.25 [Fig 1d: highest posterior density - 95% HPD = 1986.05–1995.23) using our biologically-informed priors (Table 1). Data from Stammnitz et al. [14] lead to very similar inferences (median = 1990.39, 95% HPD = 1985.85–1993.59; excluding tetraploids - median = 1991.03, 95% HPD = 1987.53–1994.08). These two independently derived estimates are quite consistent (Fig. 2) and driven by the strong temporal signal found in both datasets (using TempEst [15]; Table S3). Note that the origin date of DFTD estimated herein is more recent than that estimated in Patton et al. [2] (i.e., mid-to-late 1980s), but it remains consistent with the discovery of DFTD in 1996 and peak estimates of *R*_*E*_ ∼ 1996–1997 (Fig. 1a) when DFTD was initially discovered [16]. The excess of singletons found in [2] contributes primarily to slightly inflated terminal branch lengths, rather than affecting the overall topology of the phylogeny, and thus likely resulted in the difference in dates of origin between Stammnitz et al. [1] and Patton et al. [2].

**Table 1.**
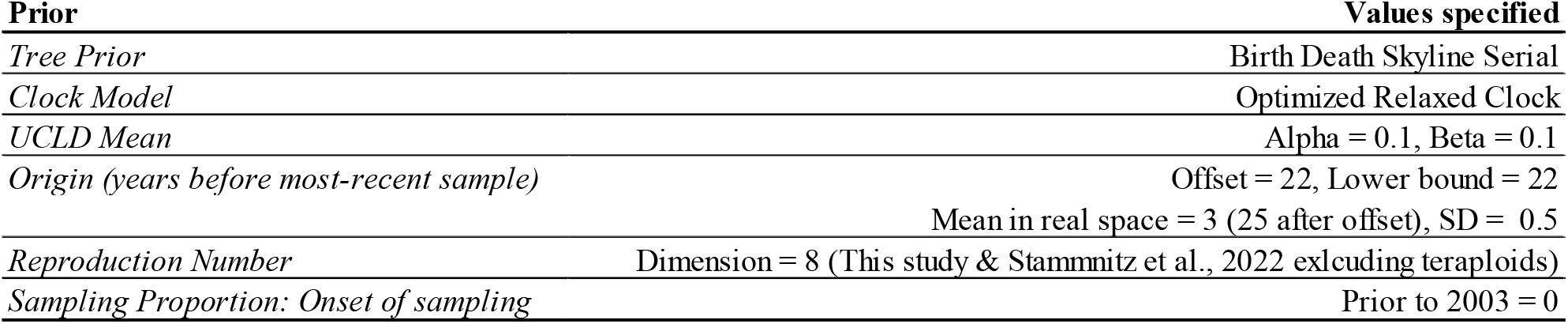
Non-default priors used in BDSKY analysis.

| **Prior** | **Values specified** |
| --- | --- |
| *Tree Prior* | Birth Death Skyline Serial |
| *Clock Model* | Optimized Relaxed Clock |
| *UCLD Mean* | Alpha = 0.1, Beta = 0.1 |
| *Origin (years before most-recent sample)* | Offset = 22, Lower bound = 22 |
|  | Mean in real space = 3 (25 after offset), SD = 0.5 |
| *Reproduction Number* | Dimension = 8 (This study & Stammnitz et al., 2022 excluding teraploids) |
| *Sampling Proportion: Onset of sampling* | Prior to 2003 = 0 |

**Figure 2.**
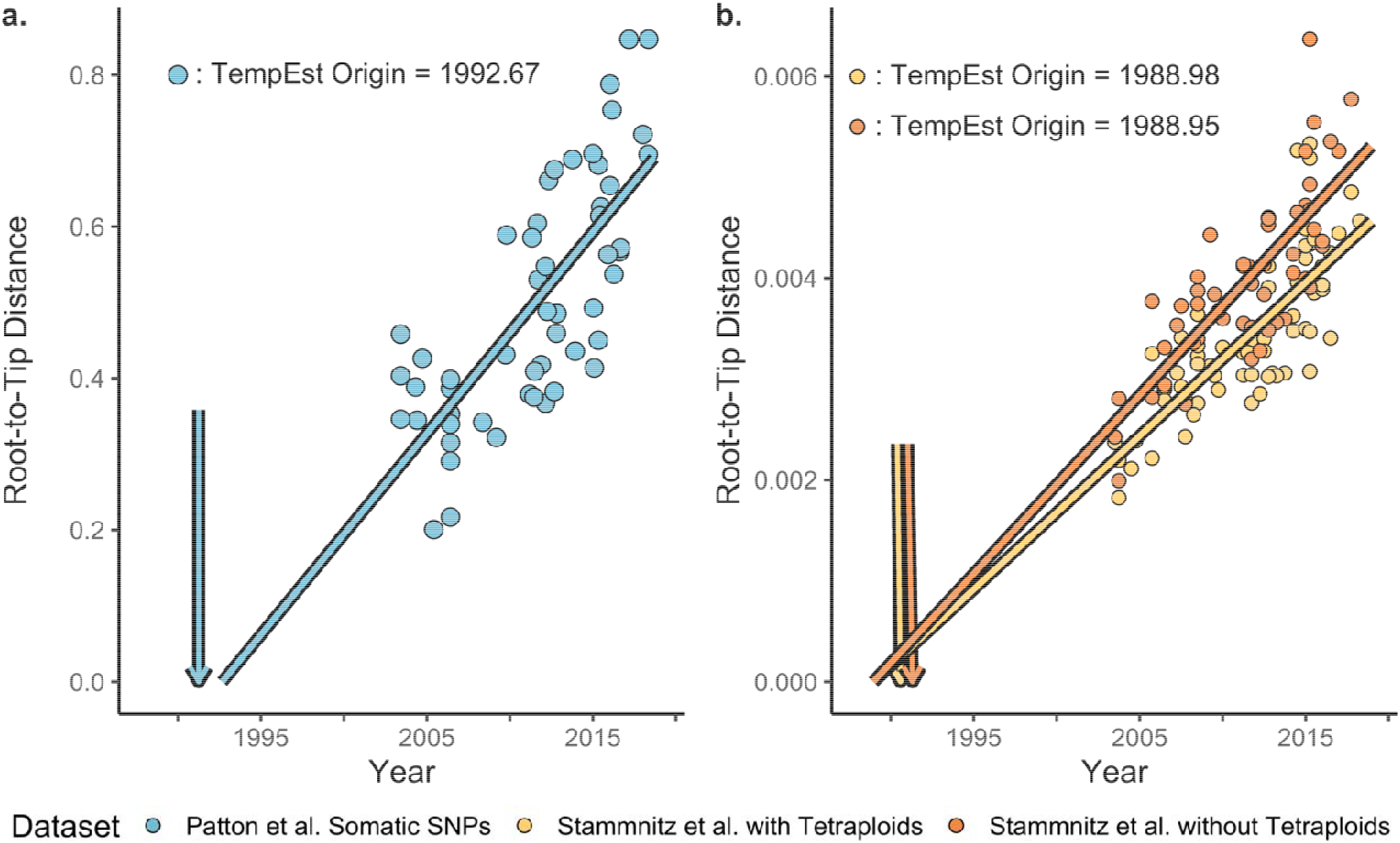
TempEst estimates of the date of DFT1 origin. TempEst applied to Maximum Likelihood phylogenies inferred using the re-filtered data from Patton et al. [2] (**a**.), or Stammnitz et al. [14] (**b**.), either including tetraploid samples. Phylogenies were re-rooted to minimize the heuristic residual mean squared error. Median estimate of DFT1 origin from BDSKY (Fig. 1d) are shown as vertical arrows.

Stammnitz et al. [1] criticize the power of our previous study [2] to detect mutations by estimating a false negative rate. However, the goal of our phylodynamic analysis was not to comprehensively identify mutations in the tumor population, but rather to estimate epidemiological parameters discussed above (i.e., directionality and rate of spread and temporal patterns in transmission rate). Stammnitz et al. [1] also assert that a minimum of 30x sequencing coverage is a general standard for genomic sequencing of cancers although in their own analysis [14], somatic mutations supported by only ≥ 3 reads were retained (Table S3). Thus, somatic mutations were called (in [14]) even if most reads (median coverage of 83x) support the reference allele, making the claim of 30x minimum coverage as necessary to call variants inconsistent with their own analyses. It is also noteworthy that the authors previously published a phylogeny of DFTD based on 1X genomic coverage (see [17]). Indeed, as is common in genomics research where sequencing costs decrease rapidly and computational speed increases, the analyses in Stammnitz et al. [1,14] include more samples with deeper sequencing coverage than our own in [2]. Lower coverage would confer a reduced ability to detect mutations relative to their study [14], but comprehensive detection of mutations across tumor genomes was not the goal of Patton et al. [2], as stated above. Nonetheless, the overall congruence between epidemiological inferences obtained using both datasets suggests that the reported differences in sequencing depth [1,14] did not meaningfully impact the inferred phylogenetic branching patterns, which are the substrate of inference of *R*_*E*_ under the Birth-death skyline model [13].

Stammnitz et al. [1] also argue that we over-estimated the mutation rate of DFTD by approximately three orders of magnitude. Note, however, that we specifically chose rapidly evolving loci to resolve phylogenies of DFTD, which is a common practice for resolving shallow evolutionary relationships (e.g., [18,19]). Consequently, our initial analyses [in 2] may have over-estimated the tumor mutation rate relative to studies with the goal of estimating genome-wide mutation rates for DFTD (as in [14]). Our revised estimates of the genome-wide substitution rate based on our refiltering strategy herein are consistent with that of a vertebrate genome (median = 2.82×10^−8^ substitutions/site/year; 95% HPD = 2.16×10^−8^–3.51×10^−8^) and within the same order of magnitude as that which we estimated from the data reported in Stammnitz et al. [1,14]: median = 8.46×10^−8^, 95% HPD = 7.45×10^−8^–9.71×10^−8^; excluding tetraploids - median = 8.39×10^−8^, 95% HPD = 7.25×10^−8^–9.57×10^−8^) (see also Table S4).

Based on the purported over-estimation of the mutation rate, Stammnitz et al. [1] argue that our resulting phylogenetic tree is incorrect and present a tree that differs substantially from the phylogenetic tree reported in Patton [2]. Their tree was inferred in IQ-Tree [20] using just 32 parsimony-informative loci copy number variants (CNVs, which are not typically used in phylogenetics or phylodynamics studies; see Table S5), which evolve in a substantially different manner than somatic point mutations used to generate our initial phylogeny [in 2]. Thus, the conflict among branching patterns in the phylogeny presented in Stammnitz et al. [1] and ours [in 2] is the result of selection of differently evolving loci to reconstruct evolutionary patterns in the two studies. Note also that Stammnitz et al. [1] collapsed CNVs to be represented as a binary state (presence/absence), consistent with that of a morphological character.

However, modelling the evolution of CNVs in this manner inaccurately represents their evolutionary process (i.e., a diploid copy number would be coded as a 0, and copy numbers of 3,4,5 or greater would all be coded as a 1), likely contributing in part to the discovery of only 32 informative sites. This likely led the CNV-based phylogeny presented in [1] to be largely composed of a polytomy, with minimal resolution of evolutionary relationships within and among three of the six clades previously described [1,14]. Further, two of their reported clades [in 14] are represented by single tumor samples and thus cannot be considered clades (nor monophyletic). Moreover, these shortcomings mean support for the branching patterns in their CNV-based phylogeny is limited and consequently cannot be used for phylodyamic epidemiological analyses (which the authors do not perform). Nonetheless, our re-analysis of both datasets supports the phylodynamic results in [2].

Stammnitz et al. [1] further argue there is evidence of substantial geographic structure among lineages of DFTD [1,14], and that the lack of geographic structure found in our data [2] relies on unrealistic migration rates of Tasmanian devils. However, evidence for substantial migration of Tasmanian devils comes from population genetics studies, showing positive genetic spatial autocorrelation extending to 120 km [21]. Moreover, the lack of geographic structure among DFTD lineages is supported by an independent population genomic analysis in [22]. Furthermore, the phylogenies in Stammnitz et al. ([14]: Fig. S3) or Kwon et al. ([17]: Fig. 1a,b) show two or three different DFTD lineages present within the most densely sampled localities and thus themselves support our finding of lack of DFTD geographic structure across Tasmania. Lastly, the fact that DFTD spread over 300 km from the east coast to the west coast of Tasmania in just over 20 years [23] supports the high migration rates necessary for lack of geographic structure among DFTD lineages.

Overall, we strongly disagree with the blanket statement [in 1] that our data are unsuitable for phylodynamic analyses and that only whole-genome data should be used for such analyses. Our study builds upon the growing body of work supporting the use of phylodynamic methods in animal systems with pathogens that evolve more slowly than viruses [12], and we see no reason that our (or their) sampling scheme should introduce pathological biases in the estimation of epidemiological parameters. We stratified our sampling with respect to geography and time specifically to avoid bias of parameter estimates for any one geographic location or time period. We contest their assertion that we sought to maximize genetic diversity, as our sample choice was blind to island-wide phylogenetic history or cladal assignment of tumors. Regardless, our re-analyses, using two different genome-wide sets of SNPs derived from *both* our data and the data used to support the critique of our work in Stammnitz et al. [1,14] clearly shows that estimates of *R*_*E*_ are robust and consistent, supporting our initial findings and conclusions [in 2] that DFTD spread is rapid and omnidirectional and that the disease is transitioning from emergence to endemism.

## 2. Materials and Methods

### 2.1 Revised data filtering procedure from Patton et al. [2]

Herein, we realigned our data from the original devil reference genome (DEVIL 7.0 [24]) to the newer chromosomal assembly reference genome, which was released after Patton et al. [2] (mSarHar1.11 [1]; Table S5) We use GATK HaplotypeCaller to jointly call of the 56 tumor samples originally analyzed in [2] according to best practices [25,26]. We stringently filtered using vcftools [27], retaining only biallelic genotypes with a mean Phred score of 60, minimum mean depth of 30x, and maximum mean depth of 100x (to account for uneven sequencing depth across samples). We then conducted our original hard-filtering step as described in [2], to prevent host genome contamination by excluding somatic mutations observed in the 12 devil genomes [5] using bcftools isec [28]. Two samples (T-146640_Wilmot_2013-07-14 & T-609040_WPP_2014-02-01) were excluded from further analysis as outliers due to unusually low counts of mutations given their sampling date. Owing to a smaller panel of host reference genomes than Stammnitz et al. [14], we further filtered to retain only SNPs present in ≥80% of our samples with a minor allele frequency of 0.05, resulting in retention of 79,397 total SNPs.

However, the minor allele frequency filter erodes temporal signal in our SNP dataset, limiting our capacity to identify rare, young, or lineage-specific alleles that increase in frequency over time. Thus, we conducted a procedure to identify clock-like somatic mutations similar to that described in [2]. Specifically, we coded the 79,397 SNPs into 012 format, wherein 0 represents the reference (Devil) allele, and 1 or 2 denotes a heterozygote or homozygote for the somatic mutation. As in [14], we recoded heterozygotes as homozygous for the somatic mutation (i.e., devil allele = 0, somatic = 1). For each SNP, we tested for a positive association of the mutation with sampling date using a logistic regression. We then excluded any SNPs with negative coefficients and ranked each SNP based on the strength of their clock-like signal. We retained the top 10% most strongly clocklike SNPs (i.e., those with the highest regression coefficients and smallest standard error), resulting in our final dataset of 3,591 SNPs (Table S1). The clock model we use relaxes the unrealistic assumption of the strict clock prior used by Stammnitz et al. [14], whereby substitution rates are assumed to be constant across all lineages and through time. These clock-like SNPs were converted to a multiple sequence alignment using the vcf2phylip python script [29].

### 2.2 Filtering procedure using data from Stammnitz et al. [1,14]

In their critique ([1]; Fig. 1B) and the supporting manuscript([14]; Fig. 1D), Stammnitz and colleagues report a time-calibrated phylogeny of DFT1 (which we refer to as “DFTD” in [2], owing to the fact that in [14], two independently evolved transmissible cancers were analyzed and referred to therein as DFT1 and DFT2). However, no phylodynamic analyses were performed in Stammnitz et al. [1] to test their assertion that there is no evidence for a transition from emergence to endemism in DFTD. Thus, we analyzed their data in a phylodynamic framework to directly test this assertion.

Using their publicly available dataset of somatic substitution and indel variants in DFT1 [14], we generated two datasets: one including and one excluding tetraploid samples (Table S3). These data report the number of reads supporting each somatic mutation as a fraction of the total reads covering the site. According to [1,14] somatic mutations were supported by ≥ 3 reads and exceeded sample-specific somatic variant allele fractions (VAF: proportion of reads supporting the mutation), or were supported by 1 read, but exceeded the sample-specific VAF of ≥ 50% of DFT1 tumors carrying the variant (Table S3). We treat read counts ≥ 3 as supporting the somatic mutation, and otherwise record the reference allele, excluding any indels. Using a custom python script, we recoded these data into a multiple sequence alignment in fasta format, excluding the hypermutated tetraploid tumor, 377T1 (denoted by the authors as ‘Clade E’), as well as the sample (102T2) attributed to the monotypic ‘Clade D’, and retained only one tumor from each devil. We then generated two datasets that either included all remaining DFT1 samples (N = 68), or excluded tetraploids (N = 53), including only those positions for which the somatic allele was present in one or more tumor samples, leading to the retention of 169,862, and 128,809 sites respectively. This leads to the retention of many singletons, and thus the clock-like signal in these data is greater than in our own. Note that our analysis of datasets including and excluding tetraploid tumor samples should recover the gross epidemiological signal present in either dataset.

We extracted approximations of sampling date from Fig. S3 in [14]. Briefly, we overlayed a vertical grid over their time-series phylogeny using their time-axis as a reference to delineate time intervals of 0.25 years. Using this grid, we rounded sampling date to the nearest quarter year in phylodynamic inference. This approach is likely to introduce noise, particularly to local branch clock rates, but in a manner that is unbiased and one that should not affect estimation of branching pattern.

### 2.3 Maximum Likelihood phylogenetic inference

Using multiple sequence alignment of both our data and the two datasets of Stammnitz et al. [14], we inferred Maximum Likelihood phylogenies using IQ-Tree v2.1.2 [20] using the GTR+G+ASC model of nucleotide substitution to account for ascertainment bias when using SNP only data. The resulting phylogenies were imported into TempEst [15] to investigate the strength of clocklike data in the Maximum Likelihood phylogeny and to identify the most probable root-location.

### 2.4 Bayesian phylodynamics analyses of both data sets

Our phylodynamic procedure using the Birth-death skyline model (BDSKY: [13]) in BEAST2 [30] is similar to the one originally described in [2]. As before, the ML phylogenies were used as starting trees. To account for the invariant sites not included in our multiple sequence alignment of clock-like SNPs, we specified the (haploid) number of each invariant base (N_A_, N_C_, N_G_, N_T_), in the reference genome to which the raw sequence data from each respective study was mapped (data from [2] mapped to Devil_ref v7.0 [30]; Data from [1,14] mapped to mSarHar1.11 [14]). We conservatively treated all sites not included in each multiple sequence alignment as invariant. We followed the procedure described here (https://groups.google.com/g/beast-users/c/QfBHMOqImFE) to modify the XML file and ‘filter’ each original alignment, accounting for the number of invariant sites. The specific constant-site counts are listed below:

**Patton:** N_A_ = 937,246,814, N_C_ =528,363,164, N_G_ = 528,188,803, N_T_ = 937,548,900

**Stammnitz (w/ Tetraploids):** N_A_ = 985,115,167, N_C_ =558,092,673, N_G_ = 557,785,141, N_T_ = 985,447,711

**Stammnitz (no Tetraploids):** N_A_ = 985,123,653, N_C_ =558,104,898, N_G_ = 557,797,061, N_T_ = 985,456,133

The site model was specified according to the substitution model inferred by IQ-Tree and all other non-default priors are found in Table 1. Using these model specifications, we ran four independent Markov Chains for a total of 300 million generations for each dataset, sampling every 10,000 generations and ran an additional Markov Chain sampling from the prior. We visually assessed that parameter estimates were not being driven by or bounded by our choice of priors, and that independent runs had converged in parameter space. Results were summarized using the bdskytools R package [31]. For our own data, all relevant parameter estimates (i.e. origin, clock rate, and estimates of *R*_*E*_) exhibited convergence across independent runs. We thus report results for the last 100 million generations from the single Markov Chain; results are qualitatively the same when summarized across the combined Markov Chains for the three other runs. For the data in [14] all four chains converged. Due to time constraints, we abbreviated these converged and well-mixed analyses at between 150 million to 200 million generations, sampling the last 100 million generations from each chain, combining them in logcombiner [29]. Similar to Stammnitz et al., [14], we corrected our estimate of DFT1’s origin by subtracting the time required for all 1,311 trunk mutations to accumulate across the posterior distribution of clock (substitution) rate estimates from the posterior distribution of disease origin.

## Supporting information

Critique and Rebutall Rejection

## Ethics

The research presented in this study was performed under University of Tasmania ethics approval A13326 and WSU IACUC approval ASAF 6796.

## Data accessibility

All data used herein are accessible on NCBI Sequence Read Archive (SRA) under BioProject PRJNA613730, BioSample Accessions SAMN14418888-906, SAMN14418908-910, SAMN14418912, SAMN14418914-931, and SAMN15869204-7. All other supporting data and code are available in the supplementary materials in Patton et al [2] and on Zenodo: https://doi.org/10.5281/zenodo.14207466; https://doi.org/10.5281/zenodo.14207593

## Declaration of AI use

We have not used AI-assisted technologies in creating this article.

## Authors contributions

ATS wrote this manuscript with contributions and edits from MJM, CPK, RH, MR-A, DGH, SC, AL, TS, HM, MJ and PAH. MJM, TS and AL assisted with data re-analysis. DGH, SC, MJ, RH, SC and MR-A provided original field samples and metadata. We thank Austin Patton for his contributions to the original manuscript and Marc Beer for assistance with re-analysis.

## Conflict of interest declarations

We declare we have no competing interests.

## Funding

This work was funded by NIH grant no. R01-GM126563 as part of the NIH-NSF-USDA Ecology and Evolution of Infectious Diseases Program (to A.S., M.E.J., H.M., and P.A.H.), by NSF DEB-2324455 to A.S., M.J.M. and R.H.), ARC DECRA grant no. DE170101116 (to R.H.).

## Notes

https://science.sciencemag.org/content/370/6522/eabb9772/suppl/DC1

https://zenodo.org/records/14207466

https://doi.org/10.5281/zenodo.14207593

### Competing Interest Statement

The authors have declared no competing interest.

