## Supplemental Table 2 for "Response to “No evidence that transmissible cancer has shifted from emergence to endemism in Tasmanian devils”"

**Table S2**

Metadata, aligned genome-wide coverage depth and purity of tumour samples from Patto

| SRA ID | Sample ID |
|---|---|
| SAMN14418888 | T-550 |
| SAMN14418889 | T-858 |
| SAMN14418890 | T-001059 |
| SAMN14418891 | T-1560 |
| SAMN14418892 | T-1773 |
| SAMN14418893 | T-2772 |
| SAMN14418894 | T-2837 |
| SAMN14418895 | T-003005 |
| SAMN14418896 | T-3115 |
| SAMN14418897 | T-3503 |
| SAMN14418898 | T-029841 |
| SAMN14418899 | T-06/1926 |
| SAMN14418900 | T-113034 |
| SAMN14418901 | T-116737 |
| SAMN14418902 | T-209336 |
| SAMN14418903 | T-209649 |
| SAMN14418904 | T-216918 |
| SAMN14418905 | T-218583 |
| SAMN14418906 | T-223262 |
| SAMN14418908 | T-495784 |
| SAMN14418909 | T-502994 |
| SAMN14418910 | T-524287 |
| SAMN14418912 | T-574450 |
| SAMN14418914 | T-584236 |
| SAMN14418915 | T-591975 |
| SAMN14418916 | T-595546 |
| SAMN14418917 | T-596114 |
| SAMN14418918 | T-608477 |
| SAMN14418919 | T-608685 |
| SAMN14418920 | T-676821 |
| SAMN14418921 | T-695064 |
| SAMN14418922 | T-699767 |
| SAMN14418923 | T-701074 |
| SAMN14418924 | T-778920 |
| SAMN14418925 | T-816986 |
| SAMN14418926 | T-826645 |
| SAMN14418927 | T-833391 |
| SAMN14418928 | T-884779 |
| SAMN14418929 | T-911814 |
| SAMN14418930 | T-918427 |

| | |
|---|---|
| SAMN14418931 | T-210100-1 |
| SAMN15869204 | T-235981 |
| SAMN15869205 | T-572899 |
| SAMN15869206 | T-574912 |
| SAMN15869207 | T-991370 |
| SAMN09242213 | T-487483; 165495-Huenc-Tumor; non-regressed (Margres et al, 2020) |
| SAMN09242220 | T-160259; Cheshite-Tumor-515; regressed (Margres et al, 2020) |
| SAMN09242222 | T-160463; Kisumu-Tumor-912; regressed (Margres et al, 2020) |
| SAMN09242223 | T-179709; Koro-Tumor-215; regressed (Margres et al, 2020) |
| SAMN09242224 | T-582387; Naya-Tumor-515; regressed (Margres et al, 2020) |
| SAMN09242226 | T-137877; Pomaire-Tumor-215; regressed (Margres et al, 2020) |

| |
|---|
| Median |

n et al, 2020.

| Location | Sampling date | Aligned coverage [x] | Purity (proportion of DFT1 DNA) |
|---|---|---|---|
| Hamilton | 01.06.2006 | 23.02 | 0.95 |
| Bothwell | 01.06.2006 | 13.93 | 0.96 |
| West Pencil Pine | 14.02.2007 | 8.59 | 0.92 |
| Sorell | 01.06.2006 | 12.43 | 0.07 |
| Bronte Park | 01.06.2004 | 17.42 | 0.83 |
| Fingal | 01.06.2006 | 11.10 | 0.95 |
| Bronte Park | 01.06.2003 | 18.39 | 0.83 |
| Franklin | 01.05.2012 | 8.67 | 0.83 |
| New Norfolk | 01.06.2005 | 37.09 | 0.47 |
| Ringarooma | 01.06.2006 | 15.11 | 0.77 |
| Narawntapu | 23.10.2009 | 13.39 | 0.55 |
| Deloraine | 01.06.2006 | 15.04 | 0.94 |
| Narawntapu | 21.09.2004 | 8.70 | 0.73 |
| Takone | 02.10.2009 | 11.59 | 0.24 |
| Lake Repulse | 22.02.2012 | 15.95 | 0.55 |
| Wynyard | 11.03.2011 | 10.42 | 0.50 |
| Warratah | 18.11.2011 | 11.17 | 0.34 |
| Black River | 11.05.2018 | 14.30 | 0.15 |
| Black River | 07.04.2017 | 7.95 | 0.11 |
| Wedge Plains | 01.09.2016 | 25.16 | 0.89 |
| Freycinet | 01.08.2011 | 5.78 | 0.76 |
| Warratah | 10.03.2009 | 9.46 | 0.91 |
| West Pencil Pine | 25.02.2016 | 19.92 | 0.26 |
| Mount William | 30.11.2015 | 36.21 | 0.51 |
| Warratah | 11.01.2016 | 15.65 | 0.13 |
| Wilmot | 06.03.2017 | 15.48 | 0.34 |
| Wilmot | 08.03.2016 | 12.05 | 0.43 |
| Elderslie | 01.11.2012 | 13.96 | 0.78 |
| Mount William | 01.12.2013 | 16.39 | 0.80 |
| Mount William | 01.09.2011 | 12.32 | 0.54 |
| Freycinet | 01.10.2013 | 13.06 | 0.26 |
| Freycinet | 04.09.2012 | 12.93 | 0.24 |
| Mount William | 19.10.2012 | 14.93 | 0.58 |
| Narawntapu | 27.04.2004 | 9.29 | 0.60 |
| Freycinet | 05.01.2018 | 21.87 | 0.91 |
| Freycinet | 23.06.2015 | 17.20 | 0.61 |
| Freycinet | 09.01.2016 | 13.58 | 0.58 |
| Narawntapu | 15.06.2011 | 17.10 | 0.57 |
| West Pencil Pine | 01.05.2018 | 12.49 | 0.17 |
| Fingal | 01.04.2012 | 14.97 | 0.89 |

| | | | |
|---|---|---|---|
| Kempton | 24.02.2012 | 9.33 | 0.53 |
| West Pencil Pine | 26.02.2015 | 12.04 | 0.52 |
| Wilmot | 01.06.2015 | 31.51 | 0.88 |
| Takone | 14.01.2015 | 18.09 | 0.80 |
| West Pencil Pine | 01.05.2011 | 83.68 | 0.88 |
| Wilmot | 01.02.2015 | 134.57 | 0.89 |
| Takone | 01.01.2015 | 63.45 | 0.15 |
| West Pencil Pine | 01.09.2012 | 57.64 | 0.69 |
| Takone | 01.01.2015 | 61.23 | 0.59 |
| Takone | 01.05.2015 | 66.31 | 0.62 |
| West Pencil Pine | 01.07.2011 | 57.90 | 0.26 |
| | | 14.97 | 0.59 |

| |
|---|
| **Aligned DFT1 coverage after accounting for purity [x]** |
| 21.83 |
| 13.40 |
| 7.89 |
| 0.87 |
| 14.45 |
| 10.55 |
| 15.19 |
| 7.15 |
| 17.45 |
| 11.56 |
| 7.41 |
| 14.21 |
| 6.35 |
| 2.79 |
| 8.70 |
| 5.21 |
| 3.81 |
| 2.08 |
| 0.85 |
| 22.34 |
| 4.40 |
| 8.57 |
| 5.22 |
| 18.54 |
| 2.01 |
| 5.25 |
| 5.14 |
| 10.87 |
| 13.17 |
| 6.62 |
| 3.33 |
| 3.09 |
| 8.66 |
| 5.54 |
| 19.80 |
| 10.51 |
| 7.90 |
| 9.77 |
| 2.15 |
| 13.40 |

| |
|---|
| 4.94 |
| 6.28 |
| 27.82 |
| 14.47 |
| 73.89 |
| 120.20 |
| 9.30 |
| 39.65 |
| 35.86 |
| 41.28 |
| 15.09 |
| 8.70 |
