## Supplemental Table 3 for "Response to “No evidence that transmissible cancer has shifted from emergence to endemism in Tasmanian devils”"

**Table S3**

DFT1 variant genotyping summary ("presence" defined as ≥3 ALT supporting

| Cohort | SRA ID | Sample ID |
|---|---|---|
| DFT1, Patton et al, 2020 | SAMN14418888 | T-550 |
| DFT1, Patton et al, 2020 | SAMN14418889 | T-858 |
| DFT1, Patton et al, 2020 | SAMN14418890 | T-001059 |
| DFT1, Patton et al, 2020 | SAMN14418891 | T-1560 |
| DFT1, Patton et al, 2020 | SAMN14418892 | T-1773 |
| DFT1, Patton et al, 2020 | SAMN14418893 | T-2772 |
| DFT1, Patton et al, 2020 | SAMN14418894 | T-2837 |
| DFT1, Patton et al, 2020 | SAMN14418895 | T-003005 |
| DFT1, Patton et al, 2020 | SAMN14418896 | T-3115 |
| DFT1, Patton et al, 2020 | SAMN14418897 | T-3503 |
| DFT1, Patton et al, 2020 | SAMN14418898 | T-029841 |
| DFT1, Patton et al, 2020 | SAMN14418899 | T-06/1926 |
| DFT1, Patton et al, 2020 | SAMN14418900 | T-113034 |
| DFT1, Patton et al, 2020 | SAMN14418901 | T-116737 |
| DFT1, Patton et al, 2020 | SAMN14418902 | T-209336 |
| DFT1, Patton et al, 2020 | SAMN14418903 | T-209649 |
| DFT1, Patton et al, 2020 | SAMN14418904 | T-216918 |
| DFT1, Patton et al, 2020 | SAMN14418905 | T-218583 |
| DFT1, Patton et al, 2020 | SAMN14418906 | T-223262 |
| DFT1, Patton et al, 2020 | SAMN14418908 | T-495784 |
| DFT1, Patton et al, 2020 | SAMN14418909 | T-502994 |
| DFT1, Patton et al, 2020 | SAMN14418910 | T-524287 |
| DFT1, Patton et al, 2020 | SAMN14418912 | T-574450 |
| DFT1, Patton et al, 2020 | SAMN14418914 | T-584236 |
| DFT1, Patton et al, 2020 | SAMN14418915 | T-591975 |
| DFT1, Patton et al, 2020 | SAMN14418916 | T-595546 |
| DFT1, Patton et al, 2020 | SAMN14418917 | T-596114 |
| DFT1, Patton et al, 2020 | SAMN14418918 | T-608477 |
| DFT1, Patton et al, 2020 | SAMN14418919 | T-608685 |
| DFT1, Patton et al, 2020 | SAMN14418920 | T-676821 |
| DFT1, Patton et al, 2020 | SAMN14418921 | T-695064 |
| DFT1, Patton et al, 2020 | SAMN14418922 | T-699767 |
| DFT1, Patton et al, 2020 | SAMN14418923 | T-701074 |
| DFT1, Patton et al, 2020 | SAMN14418924 | T-778920 |
| DFT1, Patton et al, 2020 | SAMN14418925 | T-816986 |
| DFT1, Patton et al, 2020 | SAMN14418926 | T-826645 |
| DFT1, Patton et al, 2020 | SAMN14418927 | T-833391 |
| DFT1, Patton et al, 2020 | SAMN14418928 | T-884779 |
| DFT1, Patton et al, 2020 | SAMN14418929 | T-911814 |
| DFT1, Patton et al, 2020 | SAMN14418930 | T-918427 |

| | | |
|---|---|---|
| DFT1, Patton et al, 2020 | SAMN14418931 | T-210100-1 |
| DFT1, Patton et al, 2020 | SAMN15869204 | T-235981 |
| DFT1, Patton et al, 2020 | SAMN15869205 | T-572899 |
| DFT1, Patton et al, 2020 | SAMN15869206 | T-574912 |
| DFT1, Patton et al, 2020 | SAMN15869207 | T-991370 |
| DFT1, Patton et al, 2020 | SAMN09242213 | T-487483 |
| DFT1, Patton et al, 2020 | SAMN09242220 | T-160259 |
| DFT1, Patton et al, 2020 | SAMN09242222 | T-160463 |
| DFT1, Patton et al, 2020 | SAMN09242223 | T-179709 |
| DFT1, Patton et al, 2020 | SAMN09242224 | T-582387 |
| DFT1, Patton et al, 2020 | SAMN09242226 | T-137877 |
| DFT1, Stammnitz et al, 2022 | - | 18T |
| DFT1, Stammnitz et al, 2022 | - | 32T1 |
| DFT1, Stammnitz et al, 2022 | - | 46T1 |
| DFT1, Stammnitz et al, 2022 | - | 49T1 |
| DFT1, Stammnitz et al, 2022 | - | 51T1 |
| DFT1, Stammnitz et al, 2022 | - | 52T2 |
| DFT1, Stammnitz et al, 2022 | - | 54T1 |
| DFT1, Stammnitz et al, 2022 | - | 56T2 |
| DFT1, Stammnitz et al, 2022 | - | 63T3 |
| DFT1, Stammnitz et al, 2022 | - | 78T |
| DFT1, Stammnitz et al, 2022 | - | 84T1 |
| DFT1, Stammnitz et al, 2022 | - | 86T |
| DFT1, Stammnitz et al, 2022 | - | 88T |
| DFT1, Stammnitz et al, 2022 | - | 102T2 |
| DFT1, Stammnitz et al, 2022 | - | 134T1 |
| DFT1, Stammnitz et al, 2022 | - | 134T6 |
| DFT1, Stammnitz et al, 2022 | - | 139T1 |
| DFT1, Stammnitz et al, 2022 | - | 139T4 |
| DFT1, Stammnitz et al, 2022 | - | 139T5 |
| DFT1, Stammnitz et al, 2022 | - | 139T6 |
| DFT1, Stammnitz et al, 2022 | - | 140T |
| DFT1, Stammnitz et al, 2022 | - | 141T |
| DFT1, Stammnitz et al, 2022 | - | 142T |
| DFT1, Stammnitz et al, 2022 | - | 143T |
| DFT1, Stammnitz et al, 2022 | - | 146T |
| DFT1, Stammnitz et al, 2022 | SAMN09242219 | 147Ta |
| DFT1, Stammnitz et al, 2022 | - | 155T |
| DFT1, Stammnitz et al, 2022 | SAMN09242218 | 155Ta |
| DFT1, Stammnitz et al, 2022 | SAMN15869207 | 155Tb |
| DFT1, Stammnitz et al, 2022 | - | 158T |
| DFT1, Stammnitz et al, 2022 | SAMN09242222 | 174T1a |
| DFT1, Stammnitz et al, 2022 | - | 174T2 |

| | | |
|---|---|---|
| DFT1, Stammnitz et al, 2022 | - | 199T1 |
| DFT1, Stammnitz et al, 2022 | - | 208T2 |
| DFT1, Stammnitz et al, 2022 | - | 209T3 |
| DFT1, Stammnitz et al, 2022 | - | 236T3 |
| DFT1, Stammnitz et al, 2022 | - | 341T |
| DFT1, Stammnitz et al, 2022 | - | 353T1 |
| DFT1, Stammnitz et al, 2022 | - | 356T1 |
| DFT1, Stammnitz et al, 2022 | - | 366T1 |
| DFT1, Stammnitz et al, 2022 | - | 367T1 |
| DFT1, Stammnitz et al, 2022 | - | 368T1 |
| DFT1, Stammnitz et al, 2022 | - | 372T1 |
| DFT1, Stammnitz et al, 2022 | - | 377T1 |
| DFT1, Stammnitz et al, 2022 | - | 377T3 |
| DFT1, Stammnitz et al, 2022 | - | 378T1 |
| DFT1, Stammnitz et al, 2022 | - | 379T1 |
| DFT1, Stammnitz et al, 2022 | - | 384T1 |
| DFT1, Stammnitz et al, 2022 | - | 398T1 |
| DFT1, Stammnitz et al, 2022 | - | 420T1 |
| DFT1, Stammnitz et al, 2022 | - | 421T1 |
| DFT1, Stammnitz et al, 2022 | - | 446T1 |
| DFT1, Stammnitz et al, 2022 | - | 447T1 |
| DFT1, Stammnitz et al, 2022 | - | 451T1 |
| DFT1, Stammnitz et al, 2022 | - | 458T1 |
| DFT1, Stammnitz et al, 2022 | - | 471T1 |
| DFT1, Stammnitz et al, 2022 | - | 473T3 |
| DFT1, Stammnitz et al, 2022 | - | 477T3 |
| DFT1, Stammnitz et al, 2022 | - | 528T2 |
| DFT1, Stammnitz et al, 2022 | - | 805T1 |
| DFT1, Stammnitz et al, 2022 | - | 812T2 |
| DFT1, Stammnitz et al, 2022 | - | 818T2 |
| DFT1, Stammnitz et al, 2022 | SAMN09242214 | 837T1a |
| DFT1, Stammnitz et al, 2022 | - | 876T1 |
| DFT1, Stammnitz et al, 2022 | - | 998T1 |
| DFT1, Stammnitz et al, 2022 | - | 1011T1 |
| DFT1, Stammnitz et al, 2022 | - | 1058T1 |
| DFT1, Stammnitz et al, 2022 | - | 1059T2 |
| DFT1, Stammnitz et al, 2022 | - | 1071T1 |
| DFT1, Stammnitz et al, 2022 | - | 1191T1 |
| DFT1, Stammnitz et al, 2022 | SAMN09242212 | 1439T7 |
| DFT1, Stammnitz et al, 2022 | SAMN09242213 | 2690T |
| DFT1, Stammnitz et al, 2022 | SAMN09242216 | 2691T |
| DFT1, Stammnitz et al, 2022 | SAMN09242217 | 2692T |
| DFT1, Stammnitz et al, 2022 | SAMN09242223 | 2693T |

| | | |
|---|---|---|
| DFT1, Stammnitz et al, 2022 | SAMN09242224 | 2694Ta |
| DFT1, Stammnitz et al, 2022 | SAMN09242225 | 2694Tb |
| DFT1, Stammnitz et al, 2022 | SAMN09242215 | 2695T |
| Normals, Stammnitz et al, 2022 | - | 18H |
| Normals, Stammnitz et al, 2022 | - | 32H |
| Normals, Stammnitz et al, 2022 | - | 46H |
| Normals, Stammnitz et al, 2022 | - | 51H |
| Normals, Stammnitz et al, 2022 | - | 52H |
| Normals, Stammnitz et al, 2022 | - | 54H |
| Normals, Stammnitz et al, 2022 | - | 56H |
| Normals, Stammnitz et al, 2022 | - | 63H |
| Normals, Stammnitz et al, 2022 | - | 78H |
| Normals, Stammnitz et al, 2022 | - | 84H |
| Normals, Stammnitz et al, 2022 | - | 91H |
| Normals, Stammnitz et al, 2022 | - | 134H |
| Normals, Stammnitz et al, 2022 | - | 139H |
| Normals, Stammnitz et al, 2022 | - | 141H |
| Normals, Stammnitz et al, 2022 | - | 142H |
| Normals, Stammnitz et al, 2022 | - | 143H |
| Normals, Stammnitz et al, 2022 | - | 146H |
| Normals, Stammnitz et al, 2022 | - | 147H |
| Normals, Stammnitz et al, 2022 | - | 155H |
| Normals, Stammnitz et al, 2022 | - | 158H |
| Normals, Stammnitz et al, 2022 | - | 174H |
| Normals, Stammnitz et al, 2022 | - | 199H |
| Normals, Stammnitz et al, 2022 | - | 202H |
| Normals, Stammnitz et al, 2022 | - | 203H |
| Normals, Stammnitz et al, 2022 | - | 209H |
| Normals, Stammnitz et al, 2022 | - | 212H |
| Normals, Stammnitz et al, 2022 | - | 236H |
| Normals, Stammnitz et al, 2022 | - | 338H |
| Normals, Stammnitz et al, 2022 | - | 339H |
| Normals, Stammnitz et al, 2022 | - | 340H |
| Normals, Stammnitz et al, 2022 | - | 341H |
| Normals, Stammnitz et al, 2022 | - | 353H |
| Normals, Stammnitz et al, 2022 | - | 356H |
| Normals, Stammnitz et al, 2022 | - | 366H |
| Normals, Stammnitz et al, 2022 | - | 372H |
| Normals, Stammnitz et al, 2022 | - | 377H |
| Normals, Stammnitz et al, 2022 | - | 379H |
| Normals, Stammnitz et al, 2022 | - | 398H |
| Normals, Stammnitz et al, 2022 | - | 420H |
| Normals, Stammnitz et al, 2022 | - | 421H |

| | | |
|---|---|---|
| Normals, Stammnitz et al, 2022 | - | 446H |
| Normals, Stammnitz et al, 2022 | - | 447H |
| Normals, Stammnitz et al, 2022 | - | 451H |
| Normals, Stammnitz et al, 2022 | - | 458H |
| Normals, Stammnitz et al, 2022 | - | 465H |
| Normals, Stammnitz et al, 2022 | - | 471H |
| Normals, Stammnitz et al, 2022 | - | 473H |
| Normals, Stammnitz et al, 2022 | - | 477H |
| Normals, Stammnitz et al, 2022 | - | 528H |
| Normals, Stammnitz et al, 2022 | - | 637H |
| Normals, Stammnitz et al, 2022 | - | 638H |
| Normals, Stammnitz et al, 2022 | - | 677H |
| Normals, Stammnitz et al, 2022 | - | 805H |
| Normals, Stammnitz et al, 2022 | - | 807H |
| Normals, Stammnitz et al, 2022 | - | 809H |
| Normals, Stammnitz et al, 2022 | - | 812H |
| Normals, Stammnitz et al, 2022 | - | 818H |
| Normals, Stammnitz et al, 2022 | - | 998H |
| Normals, Stammnitz et al, 2022 | - | 1011H |
| Normals, Stammnitz et al, 2022 | - | 1058H |
| Normals, Stammnitz et al, 2022 | - | 1059H |
| Normals, Stammnitz et al, 2022 | - | 1191H |
| Normals, Stammnitz et al, 2022 | - | 1334H |
| Normals, Stammnitz et al, 2022 | - | 1439H |
| Normals, Stammnitz et al, 2022 | - | 1509H |
| Normals, Stammnitz et al, 2022 | - | 1525H |
| Normals, Stammnitz et al, 2022 | - | 1528H |
| Normals, Stammnitz et al, 2022 | - | 1529H |
| Normals, Stammnitz et al, 2022 | - | 1530H |
| Normals, Stammnitz et al, 2022 | - | 1531H |
| Normals, Stammnitz et al, 2022 | - | 1532H |
| Normals, Stammnitz et al, 2022 | - | 1534H |
| Normals, Stammnitz et al, 2022 | - | 1537H |
| Normals, Stammnitz et al, 2022 | - | 1538H |
| Normals, Stammnitz et al, 2022 | - | 1545H |
| Normals, Stammnitz et al, 2022 | - | 1548H |
| Normals, Stammnitz et al, 2022 | - | 2690H |
| Normals, Stammnitz et al, 2022 | - | 2691H |
| Normals, Stammnitz et al, 2022 | - | 2693H |
| Normals, Stammnitz et al, 2022 | - | 2694H |

g reads), including truncal and clade-specific mutations, in tumour cohorts presented by Patt

| DFT1 truncal variants genotyped (% of N = 1,311) | Genotyping false negative rate (%) |
|---|---|
| 97.18 | 2.82 |
| 84.44 | 15.56 |
| 47.75 | 52.25 |
| 0.00 | 100.00 |
| 57.59 | 42.41 |
| 69.03 | 30.97 |
| 89.93 | 10.07 |
| 51.11 | 48.89 |
| 89.17 | 10.83 |
| 80.02 | 19.98 |
| 40.81 | 59.19 |
| 84.29 | 15.71 |
| 34.55 | 65.45 |
| 5.72 | 94.28 |
| 61.48 | 38.52 |
| 26.54 | 73.46 |
| 16.78 | 83.22 |
| 1.30 | 98.70 |
| 0.23 | 99.77 |
| 96.80 | 3.20 |
| 23.11 | 76.89 |
| 35.93 | 64.07 |
| 25.78 | 74.22 |
| 99.01 | 0.99 |
| 3.28 | 96.72 |
| 37.99 | 62.01 |
| 32.88 | 67.12 |
| 54.00 | 46.00 |
| 88.86 | 11.14 |
| 50.72 | 49.28 |
| 4.27 | 95.73 |
| 7.70 | 92.30 |
| 36.31 | 63.69 |
| 29.44 | 70.56 |
| 94.97 | 5.03 |
| 74.29 | 25.71 |
| 64.99 | 35.01 |
| 53.01 | 46.99 |
| 4.81 | 95.19 |
| 86.19 | 13.81 |

| | |
|---|---|
| 34.02 | 65.98 |
| 42.11 | 57.89 |
| 99.47 | 0.53 |
| 30.59 | 69.41 |
| 100.00 | 0.00 |
| 100.00 | 0.00 |
| 55.53 | 44.47 |
| 99.92 | 0.08 |
| 99.69 | 0.31 |
| 99.92 | 0.08 |
| 92.45 | 7.55 |
| 100.00 | 0.00 |
| 100.00 | 0.00 |
| 100.00 | 0.00 |
| 100.00 | 0.00 |
| 100.00 | 0.00 |
| 100.00 | 0.00 |
| 100.00 | 0.00 |
| 97.86 | 2.14 |
| 100.00 | 0.00 |
| 100.00 | 0.00 |
| 100.00 | 0.00 |
| 100.00 | 0.00 |
| 100.00 | 0.00 |
| 100.00 | 0.00 |
| 99.92 | 0.08 |
| 99.62 | 0.38 |
| 100.00 | 0.00 |
| 100.00 | 0.00 |
| 100.00 | 0.00 |
| 100.00 | 0.00 |
| 100.00 | 0.00 |
| 100.00 | 0.00 |
| 100.00 | 0.00 |
| 100.00 | 0.00 |
| 100.00 | 0.00 |
| 100.00 | 0.00 |
| 99.69 | 0.31 |
| 100.00 | 0.00 |
| 100.00 | 0.00 |
| 99.54 | 0.46 |
| 99.92 | 0.08 |
| 99.39 | 0.61 |

| | |
|---|---|
| 100.00 | 0.00 |
| 100.00 | 0.00 |
| 100.00 | 0.00 |
| 100.00 | 0.00 |
| 100.00 | 0.00 |
| 100.00 | 0.00 |
| 100.00 | 0.00 |
| 99.39 | 0.61 |
| 100.00 | 0.00 |
| 100.00 | 0.00 |
| 100.00 | 0.00 |
| 100.00 | 0.00 |
| 100.00 | 0.00 |
| 99.62 | 0.38 |
| 100.00 | 0.00 |
| 99.62 | 0.38 |
| 100.00 | 0.00 |
| 99.62 | 0.38 |
| 100.00 | 0.00 |
| 100.00 | 0.00 |
| 100.00 | 0.00 |
| 100.00 | 0.00 |
| 99.62 | 0.38 |
| 100.00 | 0.00 |
| 100.00 | 0.00 |
| 100.00 | 0.00 |
| 100.00 | 0.00 |
| 100.00 | 0.00 |
| 97.94 | 2.06 |
| 100.00 | 0.00 |
| 100.00 | 0.00 |
| 100.00 | 0.00 |
| 100.00 | 0.00 |
| 100.00 | 0.00 |
| 99.62 | 0.38 |
| 99.77 | 0.23 |
| 99.39 | 0.61 |
| 99.85 | 0.15 |
| 100.00 | 0.00 |
| 100.00 | 0.00 |
| 100.00 | 0.00 |
| 100.00 | 0.00 |
| 99.69 | 0.31 |

| | |
|---|---|
| 99.92 | 0.08 |
| 99.92 | 0.08 |
| 100.00 | 0.00 |
| 0.00 | - |
| 0.00 | - |
| 0.00 | - |
| 0.00 | - |
| 0.08 | - |
| 0.00 | - |
| 0.00 | - |
| 0.00 | - |
| 0.00 | - |
| 0.15 | - |
| 0.00 | - |
| 0.08 | - |
| 0.00 | - |
| 0.00 | - |
| 0.00 | - |
| 0.00 | - |
| 0.08 | - |
| 0.00 | - |
| 0.00 | - |
| 0.00 | - |
| 0.00 | - |
| 0.08 | - |
| 0.00 | - |
| 0.00 | - |
| 0.08 | - |
| 0.00 | - |
| 0.00 | - |
| 0.00 | - |
| 0.00 | - |
| 0.00 | - |
| 0.00 | - |
| 0.00 | - |
| 0.00 | - |
| 2.29 | - |
| 0.08 | - |
| 0.15 | - |
| 0.31 | - |
| 0.00 | - |
| 0.00 | - |
| 0.00 | - |

| | |
|---|---|
| 0.00 | - |
| 0.00 | - |
| 0.00 | - |
| 0.00 | - |
| 0.00 | - |
| 0.08 | - |
| 0.00 | - |
| 0.00 | - |
| 0.00 | - |
| 0.15 | - |
| 0.00 | - |
| 0.00 | - |
| 0.00 | - |
| 0.00 | - |
| 0.00 | - |
| 0.00 | - |
| 0.00 | - |
| 0.00 | - |
| 0.00 | - |
| 0.00 | - |
| 0.00 | - |
| 0.00 | - |
| 0.00 | - |
| 0.00 | - |
| 0.00 | - |
| 0.00 | - |
| 0.00 | - |
| 0.00 | - |
| 0.00 | - |
| 0.00 | - |
| 0.00 | - |
| 0.00 | - |
| 0.00 | - |
| 0.00 | - |
| 0.00 | - |
| 0.00 | - |
| 0.08 | - |
| 0.00 | - |
| 0.00 | - |
| 0.00 | - |

:on et al (2020) and Stammnitz et al (2023).

| DFT1 clade assignment | DFT1 clade-A1 (% of N = 328) | DFT1 clade-A2 (% of N = 139) |
|---|---|---|
| B | 0.00 | 0.00 |
| A2 | 0.00 | 82.01 |
| A2 | 0.00 | 47.48 |
| - | 0.00 | 0.00 |
| A2 | 0.00 | 56.83 |
| A2 | 0.00 | 65.47 |
| A2 | 0.00 | 84.89 |
| A2 | 0.00 | 55.40 |
| A2 | 0.00 | 85.61 |
| A2 | 0.00 | 73.38 |
| B | 0.00 | 0.00 |
| B | 0.00 | 0.00 |
| A2 | 0.00 | 28.78 |
| C | 0.00 | 0.00 |
| B | 0.00 | 0.00 |
| C | 0.00 | 0.00 |
| C | 0.00 | 0.00 |
| C | 0.00 | 0.00 |
| - | 0.00 | 0.72 |
| A2 | 0.00 | 97.12 |
| C | 0.00 | 0.00 |
| A2 | 0.00 | 25.90 |
| B | 0.00 | 0.00 |
| A2 | 0.00 | 96.40 |
| C | 0.00 | 0.00 |
| B | 0.00 | 0.00 |
| B | 0.00 | 0.00 |
| B | 0.00 | 0.00 |
| C | 0.00 | 0.00 |
| C | 0.00 | 0.00 |
| A2 | 0.00 | 4.32 |
| A1 | 3.96 | 0.00 |
| C | 0.00 | 0.00 |
| B | 0.00 | 0.00 |
| B | 0.00 | 0.00 |
| B | 0.00 | 0.00 |
| B | 0.00 | 0.00 |
| B | 0.00 | 0.00 |
| B | 0.00 | 0.00 |
| A2 | 0.00 | 89.21 |

| | | |
|---|---|---|
| B | 0.00 | 0.00 |
| A2 | 0.00 | 36.69 |
| B | 0.00 | 0.00 |
| C | 0.00 | 0.00 |
| A2 | 0.00 | 100.00 |
| B | 0.00 | 0.00 |
| C | 0.00 | 0.00 |
| C | 0.00 | 0.00 |
| C | 0.00 | 0.00 |
| C | 0.00 | 0.00 |
| A2 | 0.00 | 91.37 |
| A2 | 0.00 | 100.00 |
| B | 0.00 | 0.00 |
| C | 0.00 | 0.00 |
| A2 | 0.00 | 100.00 |
| B | 0.00 | 0.00 |
| A1 | 100.00 | 0.00 |
| A2 | 0.00 | 100.00 |
| C | 0.00 | 0.00 |
| A1 | 100.00 | 0.00 |
| A1 | 100.00 | 0.00 |
| A1 | 100.00 | 0.00 |
| B | 0.00 | 0.00 |
| A1 | 100.00 | 0.00 |
| D | 0.00 | 0.00 |
| B | 0.00 | 0.00 |
| B | 0.00 | 0.00 |
| C | 0.00 | 0.00 |
| C | 0.00 | 0.00 |
| C | 0.00 | 0.00 |
| C | 0.00 | 0.00 |
| C | 0.00 | 0.00 |
| C | 0.00 | 0.00 |
| C | 0.00 | 0.00 |
| C | 0.00 | 0.00 |
| C | 0.00 | 0.00 |
| B | 0.30 | 0.00 |
| A2 | 0.00 | 98.56 |
| A2 | 0.30 | 100.00 |
| A2 | 0.00 | 100.00 |
| A2 | 0.00 | 100.00 |
| C | 0.00 | 0.00 |
| C | 0.00 | 0.00 |

| | | |
|---|---|---|
| B | 0.00 | 0.00 |
| A2 | 0.00 | 100.00 |
| A1 | 100.00 | 0.00 |
| B | 0.00 | 0.00 |
| B | 0.00 | 0.00 |
| B | 0.00 | 0.00 |
| C | 1.22 | 0.00 |
| C | 0.00 | 0.00 |
| A2 | 0.00 | 100.00 |
| A1 | 100.00 | 0.00 |
| A2 | 0.00 | 100.00 |
| E | 0.61 | 0.00 |
| A2 | 0.00 | 100.00 |
| A2 | 0.00 | 100.00 |
| A2 | 0.00 | 100.00 |
| B | 0.00 | 0.00 |
| A2 | 0.00 | 100.00 |
| B | 0.00 | 0.00 |
| A2 | 0.00 | 100.00 |
| A2 | 0.00 | 100.00 |
| A2 | 0.00 | 100.00 |
| A2 | 0.00 | 100.00 |
| A1 | 99.39 | 0.00 |
| B | 0.00 | 0.00 |
| B | 0.30 | 0.00 |
| A2 | 0.30 | 100.00 |
| C | 0.00 | 0.00 |
| A2 | 0.00 | 100.00 |
| A1 | 99.39 | 0.00 |
| A1 | 100.00 | 0.00 |
| C | 0.00 | 0.00 |
| B | 0.00 | 0.00 |
| A2 | 0.00 | 100.00 |
| A2 | 0.00 | 100.00 |
| C | 0.00 | 0.00 |
| B | 0.00 | 0.00 |
| A2 | 0.00 | 100.00 |
| B | 0.00 | 0.00 |
| C | 0.00 | 0.00 |
| B | 0.00 | 0.00 |
| C | 0.00 | 0.00 |
| C | 0.00 | 5.04 |
| C | 0.00 | 0.00 |

| | | |
|---|---|---|
| C | 0.00 | 0.00 |
| C | 0.00 | 0.00 |
| C | 0.00 | 0.00 |
| - | 0.00 | 0.00 |
| - | 0.00 | 0.00 |
| - | 0.00 | 0.00 |
| - | 0.00 | 0.00 |
| - | 0.00 | 0.00 |
| - | 0.00 | 0.00 |
| - | 0.00 | 0.00 |
| - | 0.00 | 0.00 |
| - | 0.00 | 0.00 |
| - | 0.00 | 0.00 |
| - | 0.00 | 0.00 |
| - | 0.00 | 0.00 |
| - | 0.00 | 0.00 |
| - | 0.00 | 0.00 |
| - | 0.00 | 0.00 |
| - | 0.00 | 0.00 |
| - | 0.00 | 0.00 |
| - | 0.00 | 0.00 |
| - | 0.00 | 0.00 |
| - | 0.00 | 0.00 |
| - | 0.00 | 0.00 |
| - | 0.00 | 0.00 |
| - | 0.00 | 0.00 |
| - | 0.00 | 0.00 |
| - | 0.00 | 0.00 |
| - | 0.00 | 0.00 |
| - | 0.00 | 0.00 |
| - | 0.00 | 0.00 |
| - | 0.00 | 0.00 |
| - | 0.00 | 0.00 |
| - | 0.00 | 0.00 |
| - | 0.00 | 0.00 |
| - | 0.00 | 0.00 |
| - | 0.00 | 0.00 |
| - | 0.00 | 0.00 |
| - | 0.00 | 0.00 |
| - | 0.00 | 0.00 |
| - | 0.00 | 0.00 |
| - | 0.00 | 0.00 |

| | | |
|---|---|---|
| - | 0.30 | 0.00 |
| - | 0.00 | 0.00 |
| - | 0.00 | 0.00 |
| - | 0.00 | 0.00 |
| - | 0.00 | 0.00 |
| - | 0.00 | 0.00 |
| - | 0.00 | 0.00 |
| - | 0.00 | 0.00 |
| - | 0.00 | 0.00 |
| - | 0.61 | 0.00 |
| - | 0.00 | 0.00 |
| - | 0.00 | 0.00 |
| - | 0.00 | 0.00 |
| - | 0.00 | 0.00 |
| - | 0.00 | 0.00 |
| - | 0.00 | 0.00 |
| - | 0.00 | 0.00 |
| - | 0.00 | 0.00 |
| - | 0.00 | 0.00 |
| - | 0.00 | 0.00 |
| - | 0.00 | 0.00 |
| - | 0.00 | 0.00 |
| - | 0.00 | 0.00 |
| - | 0.00 | 0.00 |
| - | 0.00 | 0.00 |
| - | 0.00 | 0.00 |
| - | 0.00 | 0.00 |
| - | 0.00 | 0.00 |
| - | 0.00 | 0.00 |
| - | 0.00 | 0.00 |
| - | 0.00 | 0.00 |
| - | 0.00 | 0.00 |
| - | 0.00 | 0.00 |
| - | 0.00 | 0.00 |
| - | 0.00 | 0.00 |
| - | 0.00 | 0.00 |
| - | 0.00 | 0.00 |
| - | 0.00 | 0.00 |
| - | 0.00 | 0.00 |
| - | 0.00 | 0.00 |

| DFT1 clade-B (% of N = 127) | DFT1 clade-C (% of N = 349) | % DFT1 clade-D (% of N = 4,877) |
|---|---|---|
| 94.49 | 0.00 | 0.00 |
| 0.00 | 0.00 | 0.00 |
| 0.00 | 0.00 | 0.00 |
| 0.00 | 0.00 | 0.00 |
| 0.00 | 0.00 | 0.00 |
| 0.00 | 0.00 | 0.00 |
| 0.00 | 0.29 | 0.00 |
| 0.00 | 0.00 | 0.00 |
| 0.00 | 0.00 | 0.00 |
| 0.00 | 0.00 | 0.00 |
| 37.01 | 0.00 | 0.00 |
| 89.76 | 0.00 | 0.00 |
| 0.00 | 0.00 | 0.00 |
| 0.00 | 6.30 | 0.00 |
| 56.69 | 0.00 | 0.00 |
| 0.00 | 23.50 | 0.00 |
| 0.00 | 12.32 | 0.00 |
| 0.00 | 0.86 | 0.00 |
| 0.00 | 0.00 | 0.00 |
| 0.00 | 0.00 | 0.02 |
| 0.00 | 22.92 | 0.00 |
| 0.00 | 0.00 | 0.00 |
| 26.77 | 0.00 | 0.00 |
| 0.00 | 10.03 | 0.00 |
| 0.00 | 1.43 | 0.00 |
| 29.92 | 0.00 | 0.00 |
| 32.28 | 0.00 | 0.00 |
| 52.76 | 0.00 | 0.00 |
| 0.00 | 84.53 | 0.00 |
| 0.00 | 47.85 | 0.02 |
| 0.00 | 0.00 | 0.00 |
| 0.00 | 0.00 | 0.00 |
| 0.00 | 33.52 | 0.00 |
| 29.92 | 0.00 | 0.00 |
| 95.28 | 0.00 | 0.00 |
| 78.74 | 0.00 | 0.00 |
| 66.93 | 0.00 | 0.00 |
| 49.61 | 0.00 | 0.00 |
| 8.66 | 0.00 | 0.00 |
| 0.00 | 0.00 | 0.00 |

| | | |
|---|---|---|
| 29.92 | 0.00 | 0.00 |
| 0.00 | 0.00 | 0.00 |
| 100.00 | 0.00 | 0.00 |
| 0.00 | 26.65 | 0.00 |
| 0.00 | 0.00 | 0.00 |
| 100.00 | 0.00 | 0.00 |
| 0.00 | 54.15 | 0.00 |
| 0.00 | 99.43 | 0.00 |
| 0.00 | 99.71 | 0.00 |
| 0.00 | 100.00 | 0.02 |
| 0.00 | 0.00 | 0.00 |
| 0.00 | 0.00 | 0.00 |
| 100.00 | 0.00 | 0.02 |
| 0.00 | 100.00 | 0.04 |
| 0.00 | 0.00 | 0.00 |
| 100.00 | 0.00 | 0.02 |
| 0.00 | 0.00 | 0.04 |
| 0.00 | 0.00 | 0.00 |
| 0.00 | 100.00 | 0.00 |
| 0.00 | 0.00 | 0.00 |
| 0.00 | 0.00 | 0.04 |
| 0.00 | 0.00 | 0.04 |
| 100.00 | 0.00 | 0.00 |
| 0.00 | 0.00 | 0.00 |
| 0.00 | 0.00 | 99.98 |
| 100.00 | 0.00 | 0.00 |
| 100.00 | 0.00 | 0.00 |
| 0.00 | 100.00 | 0.00 |
| 0.00 | 100.00 | 0.02 |
| 0.00 | 100.00 | 0.00 |
| 0.00 | 100.00 | 0.02 |
| 0.00 | 100.00 | 0.00 |
| 0.00 | 100.00 | 0.00 |
| 0.00 | 100.00 | 0.00 |
| 0.00 | 100.00 | 0.00 |
| 0.00 | 100.00 | 0.02 |
| 100.00 | 0.00 | 0.00 |
| 0.00 | 0.00 | 0.00 |
| 0.00 | 0.29 | 0.02 |
| 0.00 | 0.00 | 0.00 |
| 0.00 | 0.00 | 0.00 |
| 0.00 | 99.71 | 0.00 |
| 0.00 | 99.43 | 0.00 |

| | | |
|---|---|---|
| 100.00 | 0.00 | 0.02 |
| 0.00 | 0.00 | 0.02 |
| 0.00 | 0.00 | 0.00 |
| 100.00 | 0.00 | 0.00 |
| 100.00 | 0.00 | 0.02 |
| 100.00 | 0.00 | 0.00 |
| 0.00 | 100.00 | 0.00 |
| 0.00 | 99.43 | 0.02 |
| 0.00 | 0.00 | 0.02 |
| 0.79 | 0.00 | 0.00 |
| 0.00 | 0.00 | 0.02 |
| 0.00 | 0.00 | 0.04 |
| 0.00 | 0.00 | 0.00 |
| 0.00 | 0.00 | 0.00 |
| 0.00 | 0.00 | 0.00 |
| 100.00 | 0.00 | 0.00 |
| 0.00 | 0.00 | 0.00 |
| 98.43 | 2.29 | 0.00 |
| 0.00 | 0.00 | 0.00 |
| 0.00 | 0.00 | 0.00 |
| 0.00 | 0.00 | 0.04 |
| 0.00 | 0.00 | 0.00 |
| 0.00 | 0.00 | 0.00 |
| 100.00 | 0.00 | 0.02 |
| 100.00 | 0.00 | 0.00 |
| 0.00 | 0.29 | 0.06 |
| 0.00 | 100.00 | 0.00 |
| 1.57 | 0.00 | 0.04 |
| 0.00 | 0.00 | 0.00 |
| 0.00 | 0.00 | 0.00 |
| 0.00 | 100.00 | 0.00 |
| 100.00 | 0.00 | 0.04 |
| 0.00 | 0.00 | 0.04 |
| 0.00 | 0.29 | 0.02 |
| 0.00 | 98.28 | 0.02 |
| 99.21 | 0.00 | 0.00 |
| 0.00 | 0.00 | 0.00 |
| 100.00 | 0.00 | 0.02 |
| 0.00 | 100.00 | 0.00 |
| 100.00 | 0.00 | 0.00 |
| 0.00 | 100.00 | 0.00 |
| 0.00 | 100.00 | 0.00 |
| 0.00 | 99.71 | 0.00 |

| | | |
|---|---|---|
| 0.00 | 100.00 | 0.02 |
| 0.00 | 100.00 | 0.00 |
| 0.00 | 100.00 | 0.00 |
| 0.00 | 0.00 | 0.02 |
| 0.00 | 0.00 | 0.02 |
| 0.00 | 0.00 | 0.04 |
| 0.00 | 0.00 | 0.00 |
| 0.00 | 0.00 | 0.00 |
| 0.00 | 0.00 | 0.00 |
| 0.00 | 0.00 | 0.00 |
| 0.00 | 0.00 | 0.00 |
| 0.00 | 0.00 | 0.02 |
| 0.00 | 0.00 | 0.00 |
| 0.00 | 0.00 | 0.00 |
| 0.00 | 0.00 | 0.00 |
| 0.00 | 0.00 | 0.00 |
| 0.00 | 0.00 | 0.00 |
| 0.00 | 0.29 | 0.00 |
| 0.00 | 0.00 | 0.00 |
| 0.00 | 0.00 | 0.00 |
| 0.00 | 0.00 | 0.00 |
| 0.00 | 0.00 | 0.00 |
| 0.00 | 0.00 | 0.00 |
| 0.00 | 0.00 | 0.00 |
| 0.00 | 0.00 | 0.02 |
| 0.00 | 0.00 | 0.00 |
| 0.00 | 0.00 | 0.00 |
| 0.00 | 0.00 | 0.00 |
| 0.00 | 0.00 | 0.00 |
| 0.00 | 0.00 | 0.02 |
| 0.00 | 0.00 | 0.00 |
| 0.00 | 0.00 | 0.00 |
| 0.00 | 0.00 | 0.00 |
| 0.00 | 0.29 | 0.00 |
| 0.00 | 0.00 | 0.00 |
| 0.00 | 0.00 | 0.00 |
| 0.00 | 4.01 | 0.02 |
| 0.00 | 0.00 | 0.00 |
| 0.00 | 0.00 | 0.02 |
| 0.00 | 0.00 | 0.00 |
| 0.00 | 0.00 | 0.00 |
| 0.00 | 0.00 | 0.00 |
| 0.00 | 0.00 | 0.00 |

| | | |
|---|---|---|
| 0.00 | 0.00 | 0.00 |
| 0.00 | 0.00 | 0.02 |
| 0.00 | 0.00 | 0.00 |
| 0.00 | 0.00 | 0.00 |
| 0.00 | 0.00 | 0.00 |
| 0.00 | 0.00 | 0.00 |
| 0.00 | 0.00 | 0.00 |
| 0.00 | 0.00 | 0.00 |
| 0.00 | 0.00 | 0.02 |
| 0.00 | 0.00 | 0.02 |
| 0.00 | 0.00 | 0.00 |
| 0.00 | 0.00 | 0.00 |
| 0.00 | 0.00 | 0.02 |
| 0.00 | 0.00 | 0.00 |
| 0.00 | 0.00 | 0.02 |
| 0.00 | 0.00 | 0.00 |
| 0.00 | 0.00 | 0.00 |
| 0.00 | 0.00 | 0.00 |
| 0.00 | 0.00 | 0.00 |
| 0.00 | 0.00 | 0.00 |
| 0.00 | 0.00 | 0.00 |
| 0.00 | 0.00 | 0.00 |
| 0.00 | 0.00 | 0.02 |
| 0.00 | 0.00 | 0.00 |
| 0.00 | 0.00 | 0.00 |
| 0.00 | 0.00 | 0.00 |
| 0.00 | 0.00 | 0.00 |
| 0.00 | 0.00 | 0.00 |
| 0.00 | 0.00 | 0.00 |
| 0.00 | 0.00 | 0.00 |
| 0.00 | 0.00 | 0.00 |
| 0.00 | 0.00 | 0.00 |
| 0.00 | 0.00 | 0.00 |
| 0.00 | 0.00 | 0.00 |
| 0.00 | 0.00 | 0.00 |
| 0.00 | 0.00 | 0.00 |
| 0.00 | 0.00 | 0.00 |
| 0.00 | 0.00 | 0.00 |
| 0.00 | 0.00 | 0.00 |
| 0.00 | 0.00 | 0.00 |

| |
|---|
| **DFT1 clade-E (% of N = 29,462)** |
| 0.00 |
| 0.00 |
| 0.00 |
| 0.00 |
| 0.00 |
| 0.00 |
| 0.00 |
| 0.00 |
| 0.00 |
| 0.00 |
| 0.00 |
| 0.00 |
| 0.00 |
| 0.00 |
| 0.00 |
| 0.00 |
| 0.00 |
| 0.00 |
| 0.00 |
| 0.00 |
| 0.00 |
| 0.00 |
| 0.00 |
| 0.00 |
| 0.00 |
| 0.00 |
| 0.00 |
| 0.00 |
| 0.00 |
| 0.00 |
| 0.00 |
| 0.00 |
| 0.00 |
| 0.00 |
| 0.00 |
| 0.00 |
| 0.00 |
| 0.00 |
| 0.00 |
| 0.00 |

| |
|---|
| 0.00 |
| 0.00 |
| 0.00 |
| 0.00 |
| 0.00 |
| 0.01 |
| 0.00 |
| 0.00 |
| 0.00 |
| 0.01 |
| 0.01 |
| 0.04 |
| 0.03 |
| 0.03 |
| 0.00 |
| 0.02 |
| 0.06 |
| 0.02 |
| 0.00 |
| 0.02 |
| 0.07 |
| 0.03 |
| 0.01 |
| 0.01 |
| 0.01 |
| 0.01 |
| 0.01 |
| 0.00 |
| 0.01 |
| 0.01 |
| 0.01 |
| 0.01 |
| 0.01 |
| 0.01 |
| 0.01 |
| 0.04 |
| 0.00 |
| 0.00 |
| 0.01 |
| 0.00 |
| 0.01 |
| 0.00 |
| 0.00 |

0.05
0.03
0.03
0.04
0.04
0.01
0.03
0.01
0.02
0.04
0.05
99.99
0.01
0.01
0.00
0.00
0.04
0.00
0.01
0.02
0.05
0.01
0.01
0.04
0.05
0.05
0.01
0.03
0.01
0.01
0.00
0.01
0.03
0.01
0.01
0.00
0.00
0.01
0.00
0.01
0.01
0.00
0.00

| |
|---|
| 0.01 |
| 0.00 |
| 0.01 |
| 0.01 |
| 0.01 |
| 0.02 |
| 0.00 |
| 0.01 |
| 0.00 |
| 0.00 |
| 0.01 |
| 0.01 |
| 0.00 |
| 0.01 |
| 0.00 |
| 0.01 |
| 0.00 |
| 0.00 |
| 0.01 |
| 0.03 |
| 0.00 |
| 0.00 |
| 0.00 |
| 0.00 |
| 0.02 |
| 0.00 |
| 0.00 |
| 0.01 |
| 0.00 |
| 0.01 |
| 0.00 |
| 0.00 |
| 0.00 |
| 0.03 |
| 0.00 |
| 0.00 |
| 0.00 |
| 0.02 |
| 0.01 |
| 0.00 |
| 0.01 |
| 0.01 |
| 0.01 |

| |
|---|
| 0.00 |
| 0.00 |
| 0.00 |
| 0.00 |
| 0.00 |
| 0.01 |
| 0.02 |
| 0.02 |
| 0.01 |
| 0.00 |
| 0.00 |
| 0.00 |
| 0.01 |
| 0.00 |
| 0.00 |
| 0.01 |
| 0.00 |
| 0.00 |
| 0.01 |
| 0.00 |
| 0.00 |
| 0.00 |
| 0.00 |
| 0.00 |
| 0.01 |
| 0.01 |
| 0.01 |
| 0.00 |
| 0.01 |
| 0.00 |
| 0.00 |
| 0.02 |
| 0.01 |
| 0.01 |
| 0.00 |
| 0.00 |
| 0.00 |
| 0.01 |
| 0.01 |
| 0.00 |
