## Supplemental Table 4 for "Response to “No evidence that transmissible cancer has shifted from emergence to endemism in Tasmanian devils”"

**Table S5**

Somatic substitutions identified within the subset of all "clock-like genes" (Pa

| ENSEMBL ID (v105) | HGNC | DEVIL7.0 Scaffold ID |
|---|---|---|
| ENSSHAG00000002874 | ARL14EPL | GL842139 |
| ENSSHAG00000003887 | MYO1F | GL841425 |
| ENSSHAG00000023180 | GDF6 | GL841368 |
| ENSSHAG00000001479 | | GL842771 |
| ENSSHAG00000003540 | Zinc finger protein 665-like | GL835033 |
| ENSSHAG00000001804 | TRHR2 | GL834768 |
| ENSSHAG00000009187 | GKN1 | GL834702 |
| ENSSHAG00000016343 | ANKRD60 | GL834665 |
| ENSSHAG00000010978 | INPP5F | GL834460 |
| ENSSHAG00000011106 | MRPS22 | GL849592 |
| ENSSHAG00000000980 | TMEM45B | GL849688 |
| ENSSHAG00000011332 | BCO2 | GL849661 |
| ENSSHAG00000002177 | TRIM25 | GL857553 |
| ENSSHAG00000003546 | INTS12 | GL864760 |
| ENSSHAG00000003826 | CAPN6 | GL867616 |

tton et al, 2020) matched between the DEVIL7.0 (2012) and mSarHar1.11 (2022) reference ge

| DEVIL7.1 Scaffold ID | DEVIL7.1 Start | DEVIL7.1 End | mSarHar1.11 Chromosome |
|---|---|---|---|
| Chr2_supercontig_000000998 | 44222 | 53237 | 1 |
| Chr2_supercontig_000000284 | 78972 | 111992 | 1 |
| Chr2_supercontig_000000227 | 15 | 24826 | 1 |
| Chr2_supercontig_000001630 | 11834 | 14490 | 1 |
| Chr1_supercontig_000000621 | 66434 | 68008 | 2 |
| Chr1_supercontig_000000356 | 17854 | 39972 | 2 |
| Chr1_supercontig_000000290 | 514136 | 521595 | 2 |
| Chr1_supercontig_000000253 | 2195465 | 2203951 | 2 |
| Chr1_supercontig_000000048 | 800979 | 825590 | 2 |
| Chr3_supercontig_000000070 | 797767 | 816133 | 3 |
| Chr3_supercontig_000000166 | 5440 | 27384 | 3 |
| Chr3_supercontig_000000139 | 833339 | 871968 | 3 |
| Chr4_supercontig_000000834 | 25362 | 57064 | 4 |
| Chr6_supercontig_000000036 | 66708 | 98568 | 6 |
| Chrx_supercontig_000000048 | 76706 | 91165 | X |

enomes.

| mSarHar1.11 Start | mSarHar1.11 End |
|---|---|
| 234459134 | 234469763 |
| 672392632 | 672492067 |
| 576854300 | 576876516 |
| 714683179 | 714693224 |
| 34278710 | 34292399 |
| 47558285 | 47580465 |
| 114821840 | 114829309 |
| 235191998 | 235279890 |
| 577183100 | 577359768 |
| 312026829 | 312061184 |
| 454158622 | 454221787 |
| 488312724 | 488350315 |
| 347426155 | 347459015 |
| 133126598 | 133164670 |
| 77342222 | 77356616 |

| Somatic DFT1 substitutions, identified across 78 tumour genomes (Stammnitz et al, 2022) |
|---|
| 1 |
| 5 |
| 2 |
| 1 |
| 1 |
| 0 |
| 0 |
| 1 |
| 6 |
| 0 |
| 1 |
| 1 |
| 1 |
| 1 |
| 1 |
