## Supplementary material for "Response to “No evidence that transmissible cancer has shifted from emergence to endemism in Tasmanian devils”": Critique and Rebutall Rejection

Home | New Submission () | My Account

(https://www.science.org/journals)

**Journal:** Science
**Manuscript ID:** abq7783

**Status:** Rejected

**Article Type:** Technical Response

**First Author:** Austin Patton

**Assigned To:** Sacha Vignieri

MS Info () | Authors () | MS Documents () | Dataset ()

**Journal:** Science

**Article Type:** Technical Response

**Assigned To:** Sacha Vignieri

**Title:** Response to “No evidence that a transmissible cancer has shifted from emergence to endemism in Tasmanian devils”

**Short Title:**

**Field Codes:**

**Primary**

*There are no Field Codes selected*

**Secondary**

*There are no Field Codes selected*

Home | New Submission () | My Account

Welcome Andrew Logout

(https://www.science.org/journals)

**Abstract:**

The title of the technical comment, “No evidence that a transmissible cancer has shifted from emergence to endemism in Tasmanian devils” (1) is misleading. That is, the authors fail to provide any epidemiological evidence contradicting the findings of Patton et al. (2), and ignore decades of independent field data, genomic data, and models that support the declining threat of devil facial tumor disease (3–9). Stammnitz et al. (1) correctly draw attention to the fact that our data filtering procedures could have been more stringent in Patton et al. (2), and accordingly, we re-analyze our data, as well as the data provided in their supporting manuscript (10). Phylodynamic re-analyses of our data and independent analysis of their data both support the findings of Patton et al. (2); the effective reproductive number of the transmissible cancer gradually declines to one at present. These data are consistent with the original conclusion (2) that DFTD is transitioning from emergence to endemism.

**Teaser:**

**Funding Source:**

AAAS

© 2025 American Association for the Advancement of Science. All Rights Reserved. AAAS is a partner of HINARI, AGORA, OARE, ORCID, CrossRef, and COUNTER.

Privacy Policy (https://www.science.org/content/page/privacy-policy) | Submission Help ()
